# The genetic basis of wing spots in *Pieris canidia* butterflies

**DOI:** 10.1101/2022.11.17.516903

**Authors:** Jocelyn Liang Qi Wee, Suriya Narayanan Murugesan, Christopher Wheat, Antónia Monteiro

**Author notes:** Joint first authors (Jocelyn Liang Qi Wee, Suriya Narayanan Murugesan).

## Abstract

Spots in pierid butterflies and eyespots in nymphalid butterflies are likely non-homologous wing colour pattern elements, yet they share a few features in common. Both develop black scales that depend on the function of the gene *spalt*, and both might have central signalling cells. This suggests that both pattern elements may be sharing common genetic circuitry. Hundreds of genes have already been associated with the development of nymphalid butterfly eyespot patterns, but the genetic basis of the simpler spot patterns on the wings of pierid butterflies has not been investigated. To facilitate studies of pierid wing patterns, we report a high-quality draft genome assembly for *Pieris canidia*, the Indian cabbage white. We then conducted transcriptomic analyses of pupal wing tissues sampled from the spot and non-spot regions of *P. canidia* at 3-6h post-pupation. A total of 1352 genes were differentially regulated between wing tissues with and without the black spot, including *spalt, Krüppel-like factor 10*, genes from the Toll, Notch, TGF-β, and FGFR signalling pathways, and several genes involved in the melanin biosynthetic pathway. We identified 21 genes that are up-regulated in both pierid spots and nymphalid eyespots and propose that spots and eyespots share regulatory modules despite their likely independent origins.

## Introduction

Discovering how novel traits originate and diversify across different lineages of organisms is a long-standing pursuit of evolutionary biologists [1]. Most morphological traits are the result of complex gene-regulatory networks (GRNs), orchestrated by intercellular signals and their target transcription factors (TFs). The level of complexity underlying the development of any trait is usually so high that it has been proposed that novel traits originate via the rewiring of partial or whole pre-existent GRNs into new developmental contexts [2]. An example involves the gene *optix*, a gene essential for eye development which appears to have been redeployed, together with many other eye pigment enzyme genes, onto butterflies’ wings to differentiate red wing patterns [3-6]. Another example involves the co-option of a general limb GRN, regulating the development of antennae, legs, and wings, onto a small group of cells on the wing to underlie the evolution of nymphalid eyespot patterns [7].

Eyespot patterns, however, appear to have replaced pre-existing simpler spots that were already present on the margins of nymphalid wings. Eyespot and spot patterns could, thus, potentially share common positional information during development conferred by a common set of genes. This evolutionary transition, of spots to eyespots, inferred from the use of a large phylogeny of the butterfly family Nymphalidae and comparative methods [8], has not yet been fully explored at the genetic and developmental levels. A recent study, however, that examined the function of two essential genes for the differentiation of eyespot centers, *Distal-less*, and *spalt*, found that they played no such role in differentiating the centers of spot patterns in a separate family of butterflies, the Pieridae, which last shared a common ancestor with Nymphalidae nearly 100 million years ago [9]. This study suggested, instead, that spots in pierids are likely homologous to more marginal parafocal elements found on the wings of butterflies belonging to the family Nymphalidae, and are not positional homologs of spot patterns in nymphalid lineages that eyespots are thought to have evolved from [10].

Yet, pierid spots and eyespots of *Bicyclus anynana* butterflies have a few features in common that might signal process homology at the level of the constituent gene-regulatory networks (GRNs) [11, 12]. They both have black scales that require the gene *spalt* for their differentiation [7, 10]; and both patterns become reduced in size when cells at their center are damaged in the early pupal stage [13, 14]. The full extent of shared genes, and larger GRNs, between spots in pierids and eyespots in nymphalids, however, is unknown.

To further probe the genetic basis of spots, and to compare it with that of eyespots, we first assembled and annotated a high-quality draft genome for *P. canidia*, generated from long reads obtained from the PacBio Sequel platform. RNA-seq was performed to identify the suite of transcription factors and signalling pathway molecules that underlie the development of spot patterns in the wings of *Pieris canidia*, following the same protocol used to probe the genetic basis of nymphalid eyespots [15]. This involved micro-dissections of very small pieces of wing tissue containing the spot region, and of flanking tissue without spots during the early pupal stage, followed by sequencing and differential gene expression analysis. We then compared the gene expression profiles involved in the development of both spots and eyespots in butterflies.

## Results

### *P. canidia* genome assembly and annotation

The genome profile of *P. canidia* was determined using a 19-kmer distribution from Illumina short reads from single individual using GenomeScope (http://qb.cshl.edu/genomescope/) [16]. The k-mer counting was done using Jellyfish (v.2.2.3) [17] and the approximate genome size was determined to be around 224 MB (Figure S1). We assembled the PacBio long reads into contigs using wtdbg2 (version2.4) [18] and canu (version1.9) [19] assembler separately. Canu produced an assembly of 325 MB size with N50 length of 767 KB. Wtdbg2 produced a 257 MB assembly with a N50 length of 2.7 MB. The assemblies from both assemblers were merged using quickmerge [20] followed by purging of the haplotigs to remove heterozygosity using purge_haplotigs [21]. Subsequent output from purge_haplotigs was polished using long reads and short reads with three rounds of racon [22] and pilon [23] respectively, resulting in a final assembly of 264.9 MB size and a N50 length of 10.1 MB with 392 contigs (**Table 1**). The assembly was repeat masked using RepeatMasker [24] and annotated with four rounds of Maker [25] using the transcriptome constructed from RNA-seq from different tissues (detailed in Materials and Methods section) and transcriptome and proteins sequences from other species of butterflies as relative species information. This resulted in 14518 genes with 26376 transcripts. We checked the completeness of the assembly and gene set using a lepidoptera database with Benchmarking Universal Single-Copy Orthologs (BUSCO) (last accessed October 2022) [26]. The genome was very complete with 97.5% of single copy gene sets found in the *P. canidia* genome (**Table 1**).

**Table 1.**
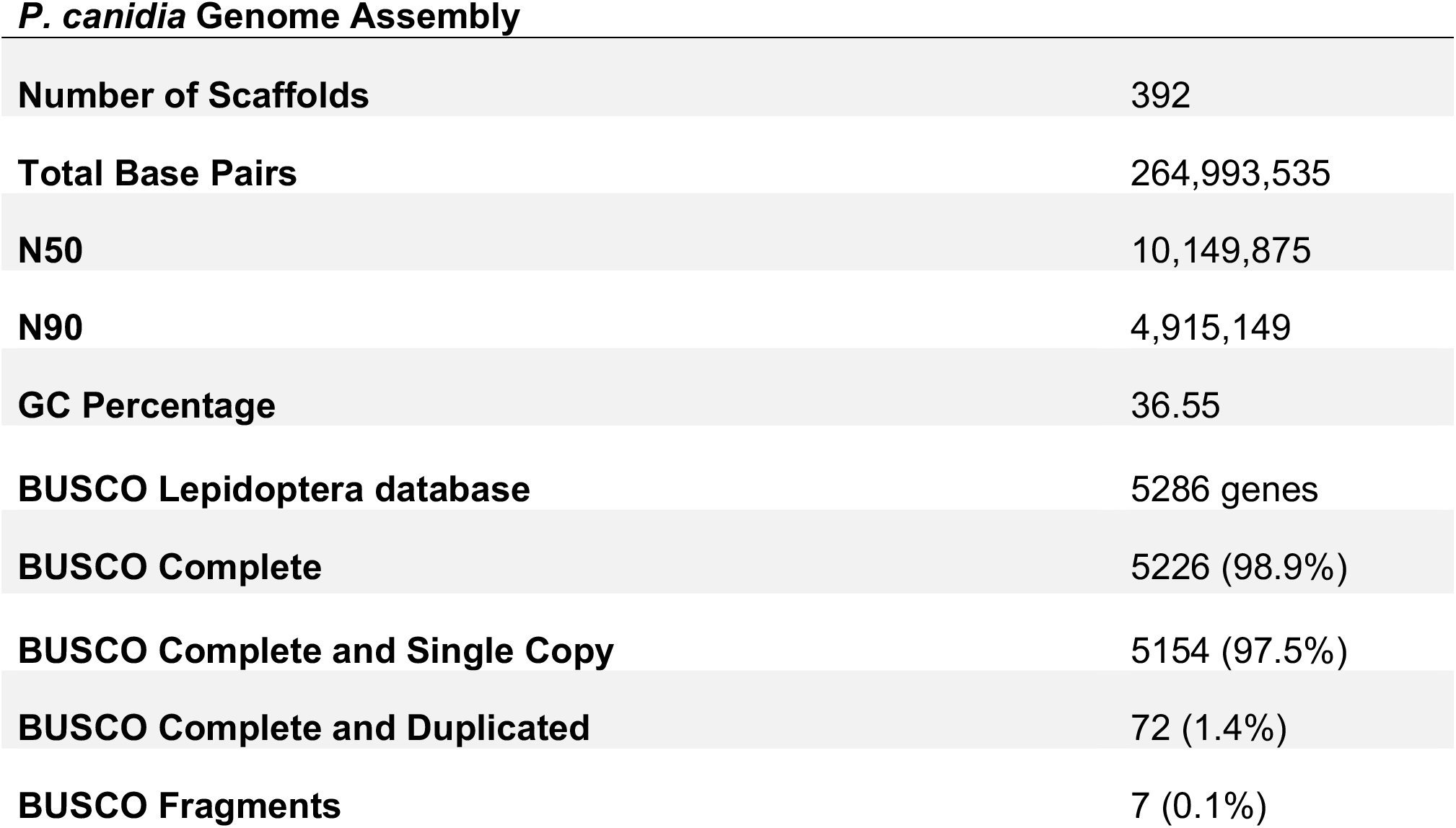
Genome assembly statistics of *P. canidia*.

### Transcriptomic analysis performed on regions of spot patterns of *P. canidia*

To identify the genes and signalling pathways that are responsible for organising the spot patterns of *P. canidia*, we compared gene expression patterns between regions of pupal forewing tissues that will later develop spots (spot region in M3) versus two regions of the wings that do not develop spots (the proximal region next to the spot in the same M3 sector and the Cu1 region, in a more posterior sector) (Fig 1A & 1B).

**Figure 1.**
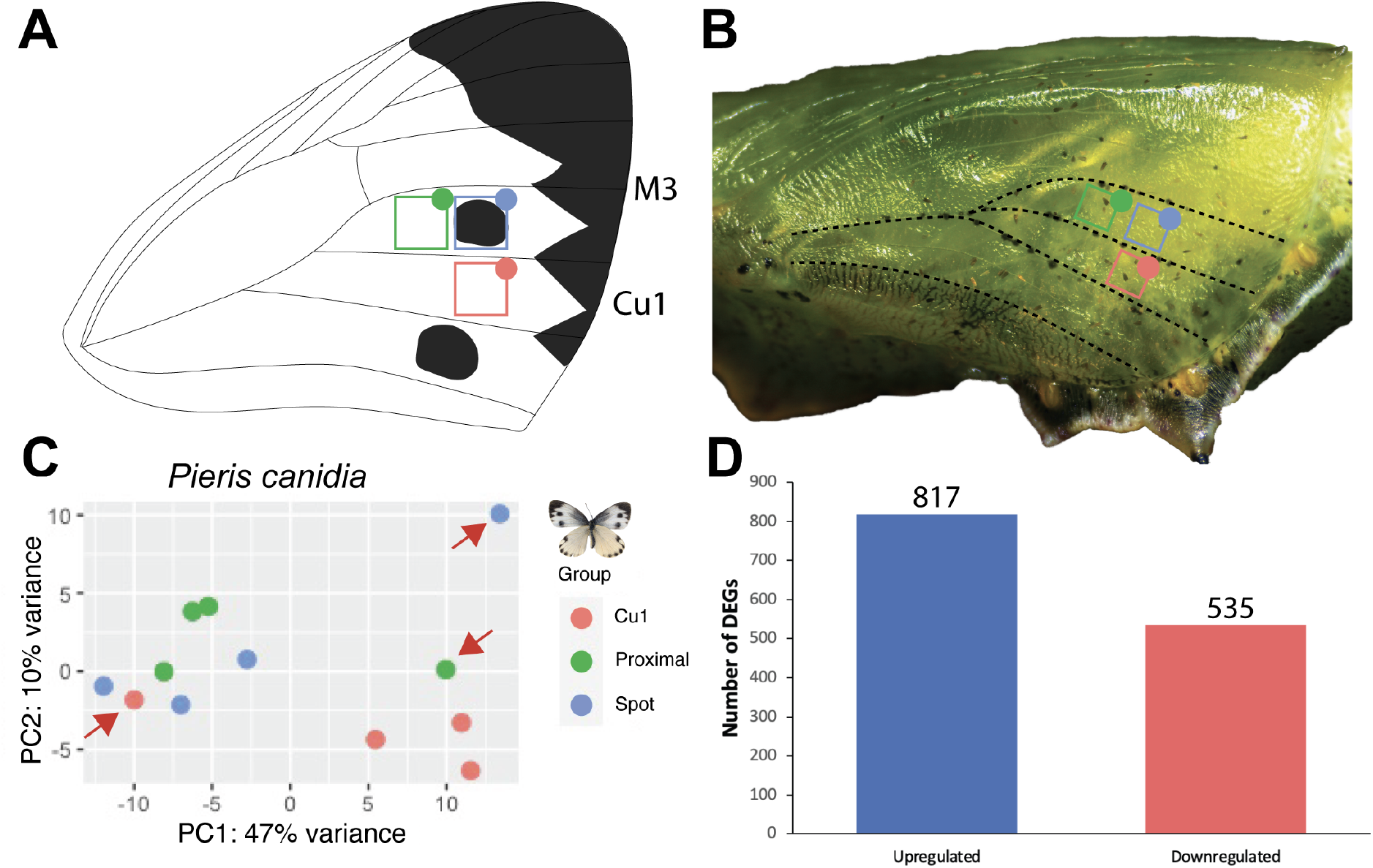
Data analyses of RNA-seq samples. A) Drawing of a *P. canidia* adult forewing showing the approximate location of the three dissected areas mapped to the wing. B) *P. canidia* pupa (3h old) with the three dissected tissues marked on the wing. C) Principal component analyses of RNA samples clustered by the area of the wing of which samples were dissected. Apart from three outlier groups, the rest of the samples clustered first with others from the same wing location. (Red: Wing tissues dissected from the Cu1 wing sector, Green: Wing tissues dissected from an area that is proximal to the spot pattern, and Blue: Wing tissues dissected from an area that contains the spot pattern. D) Number of differentially expressed genes (DEGs) for the comparison of M3 “Spot” versus Cu1 “Control” tissue.

Wing tissues were dissected at 3-6 hrs after pupation, which was the time window previously sampled for eyespot tissue in *Bicyclus anynana* [7, 15]. This period is when the central focal cells are beginning the process of colour ring differentiation in nymphalid eyespots, because when these cells are transplanted elsewhere on the wing, they can set-up rings of colour around them [13, 27]. In total, 12 libraries were sequenced (4 biological reps/sample type) with each replicate sequenced to a depth of between 40-90 million reads on average.

Principal component analysis (PCA) of the libraries revealed that the samples generally clustered together based on the location of the wing where the tissue was obtained, except for 1 library from each group (Fig 1C). Samples from the same wing sector clustered together, i.e., the group ‘spot’ clustered closely together with the group ‘proximal’, whereas tissues belonging to a different wing sector, ‘Cu1’ clustered separately. Since no morphological marker distinguished the two wing locations within the M3 sector, it is possible that the two dissected tissues might have overlapped the spot region in some cases. As the PCA plot revealed 3 outliers that deviated significantly from their respective groups, these were removed for downstream analyses. Subsequent hierarchical clustering of samples showed that samples belonging to the same group clustered together (Fig S2).

### Genes differentially expressed in the wing spot pattern

Two comparisons were performed to identify differentially expressed genes (DEGs) in spots of *P. canidia*: we compared ‘spot’ vs ‘proximal’ and ‘spot’ vs ‘cu1’ to obtain the common set of genes that appeared in both comparisons. The comparison of ‘spot’ vs ‘proximal’ yielded 15 upregulated genes and 15 downregulated genes in spots whereas the comparison of ‘spot’ vs ‘cu1’ tissues yielded 817 upregulated genes and 535 downregulated genes in spots (Fig 1D). Overall, 16 DEGs were found in common across both comparisons with 4 genes being differentially regulated in a conflicting manner between the two comparisons (Table S1). Of these 16 genes, 4 genes were not annotated or were identified as uncharacterised proteins. The relatively low number of DEGs found in common across both sets of comparisons was primarily constrained by the low number of DEGs between ‘spot’ and ‘proximal’ tissues. Given that variations in spot size and location exist in natural populations, the possibility that wing tissue encompassing the spot pattern may have overlapped between the two tissue groups dissected side-by-side from the same wing sector, cannot be ruled out. By focusing solely on these 16 DE annotated genes, there is a risk of missing out on additional candidate genes and pathways responsible for organising the spot pattern.

Thus, to better elucidate the full suite of genes that are associated with spot development, we decided to focus on the 1352 DEGs, with a p-value adjusted (padj) of <0.01, that are up- or downregulated between ‘spot’ and ‘cu1’ tissues alone (Fig 2). Upregulated DEGs were enriched for several GO terms such as “structural constituent of cuticle”, “structural constituent of chitin-based cuticle”, “chitin-based extracellular matrix”, and other GO terms. Downregulated DEGs were enriched for GO functions relating to “defense response to bacterium”, “regulation of transcription, DNA-templated”, and “protein phosphorylation” (Fig S3). Consistent with previous findings, *spalt* (*sal*) (Log_2_FC: 1.05) is included in the set of DEGs and is shown to be over-expressed in wing tissues containing the spot pattern. This shows that our chosen method of analysis and thresholds applied to the dataset is sensitive enough to detect genes that are involved in wing spot development.

**Figure 2.**
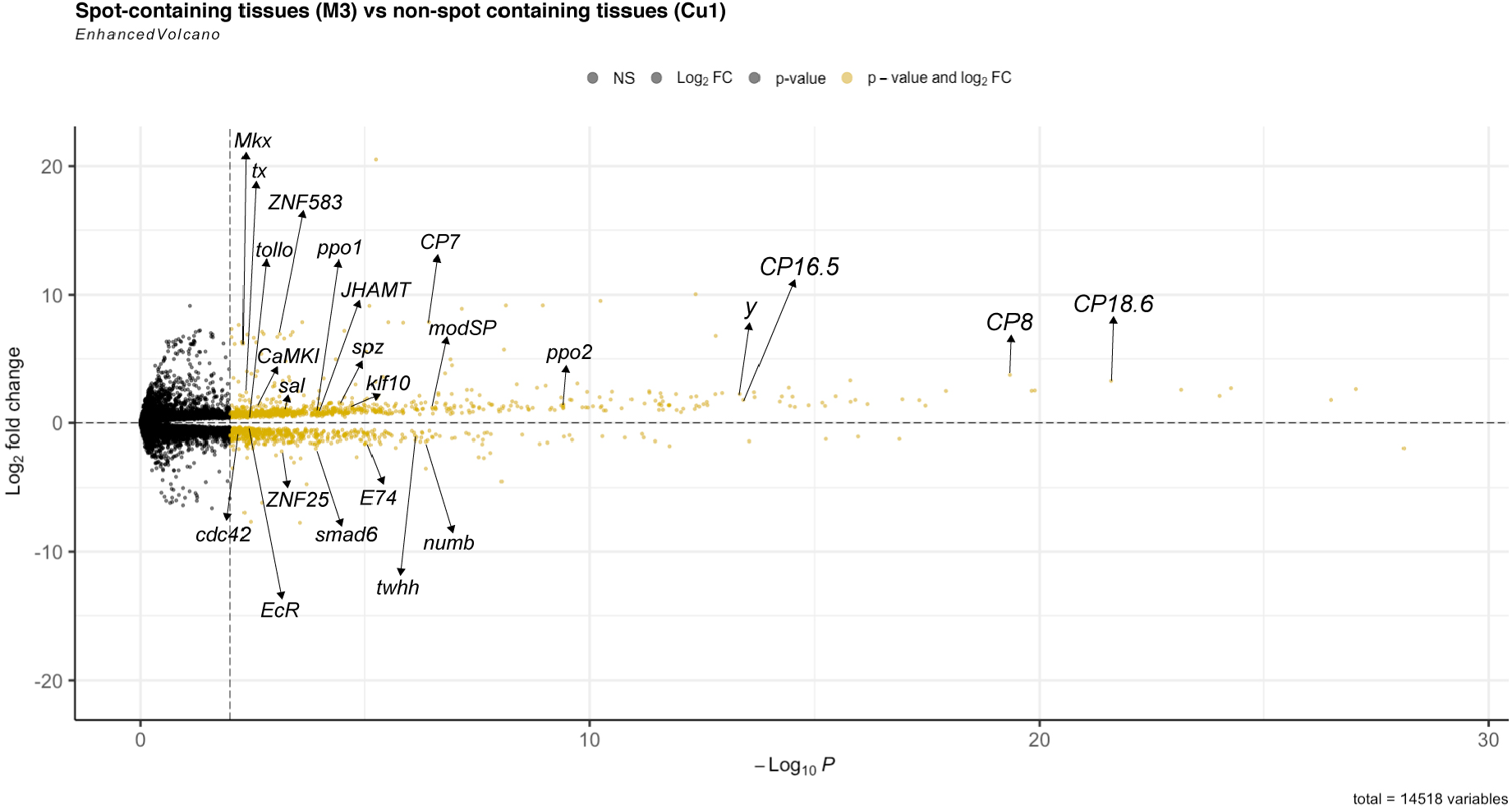
Volcano plot of DEGs between “spot” and “cu1” tissues of 3-6h *P. canidia* pupal forewings. Log_2_-fold change values were plotted against average values of -log_10_(p-value) with the threshold of DEGs set at padj <0.01. DEGs are marked as yellow dots. Genes upregulated in spot tissue are on the top half of the volcano plot. Downregulated genes are on the bottom half of the volcano plot. This plot is generated using the R package EnhancedVolcano [28].

### Pigmentation genes associated with spots

As the wing spot pattern of *P. canidia* consists of black scales, we looked through the list of DEGs to identify those known to function in pigment and melanin biosynthesis. In total, 10 out of 1352 DEGs are pigment-associated genes (Table 2). Of these 10 genes, 9 are involved in the melanogenic pathway in insects, and 1 in the pteridine biosynthesis pathway. All 10 genes involved in pigmentation processes are significantly upregulated in spot tissue compared to Cu1 control tissue, with *protein yellow-like* showing the largest fold change of 2.24 respectively. No genes involved in the melanin biosynthetic pathway or any other pigment pathways (i.e., ommochrome/pteridine) were downregulated at this stage of development.

**Table 2.** Genes annotated as part of pigment producing pathways expressed in spot tissues. Their log_2_FC, padj values, and annotated biological processes. All genes described in this table are those that are upregulated in Spot vs Cu1 samples.

| Gene ID | Gene Name | Log<sub>2</sub>FC | padj | Biological Processes |
| --- | --- | --- | --- | --- |
| Pcan_01059 | *phenoloxidase subunit 1* | 0.9761 | 0.0001 | melanin biosynthetic process from tyrosine |
| Pcan_04990 | *phenoloxidase subunit 2-like* | 1.1925 | 3.91E-10 | melanin biosynthetic process from tyrosine |
| Pcan_06980 | *protein yellow-like* | 1.5657 | 5.74E-05 | melanin biosynthetic process |
| Pcan_06984 | *major royal jelly protein 1-like isoform X2* | 0.6703 | 0.0003 | melanin biosynthetic process |
| Pcan_07144 | *pyrimidodiazepine synthase-like* | 0.7874 | 0.0034 | eye pigment biosynthetic process; pteridine biosynthetic process |
| Pcan_09922 | *L-dopachrome tautomerase yellow-f-like* | 0.9512 | 0.0003 | melanin biosynthetic process from tyrosine |
| Pcan_10393 | *protein yellow-like* | 2.2439 | 4.59E-14 | melanin biosynthetic process from tyrosine |
| Pcan_11552 | *protein yellow* | 1.0028 | 0.0001 | melanin biosynthetic process |
| Pcan_11721 | *cysteine sulfinic acid decarboxylase* | 1.0397 | 0.0071 | developmental pigmentation |
| Pcan_12794 | *protein yellow-like* | 1.1784 | 3.40E-09 | melanin biosynthetic process |

### Transcription factors and signalling pathways

The re-wiring of pre-existing gene regulatory networks to novel contexts has been proposed to drive the evolution of novel traits. Central to this regulation are ligands, receptors, and transcription factors involved in signalling pathways. To better understand the regulatory landscape that is controlling the expression of effector genes that are responsible for producing the dark wing spot, regulatory genes were identified by GO analysis. This was achieved through searching through the set of annotated DE genes with corresponding GO terms using keywords such as i) “DNA-binding transcription factor activity” (GO:0003700, GO0000981, GO0001228, GO:0001217), (ii) “signalling pathway” and iii) “signal transduction”. Sixteen transcription factors were differentially expressed between the two tissues (Spot vs Cu1) (Table 3). Unlike the pigment pathway genes, most of the transcription factors were significantly downregulated in spots as summarised in Table 3. Other transcription factors, such as *Krüppel-like factor 10, homeotic protein spalt-major-like isoform X1*, and *helix-loop-helix protein delilah-like* were upregulated in spot tissues.

**Table 3.**
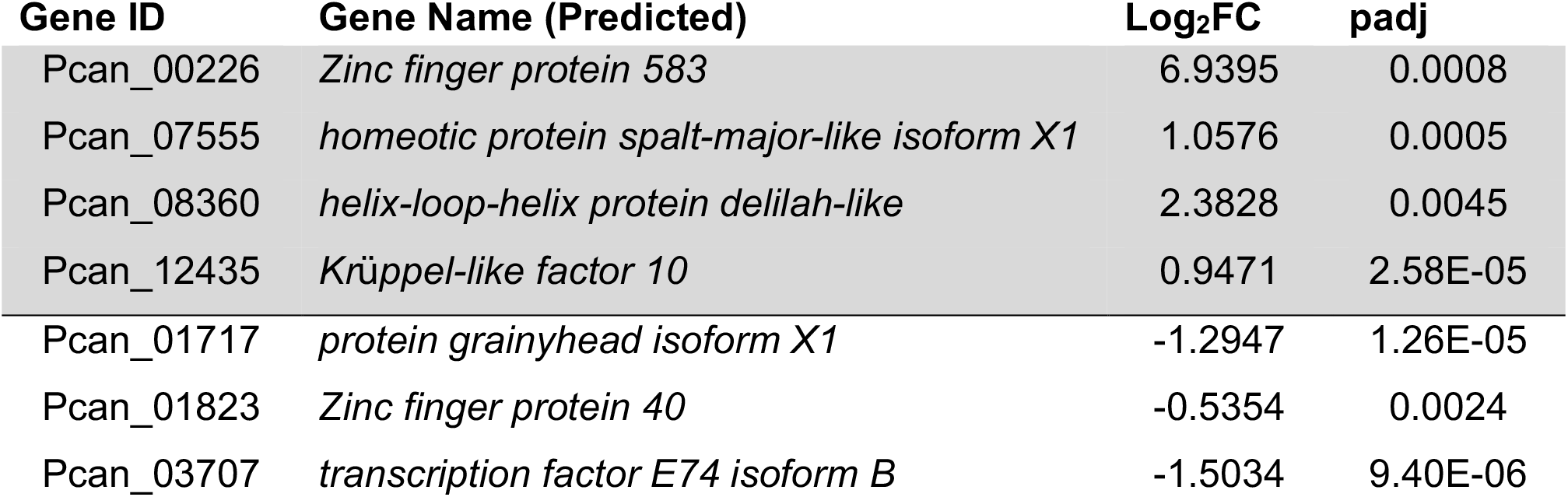
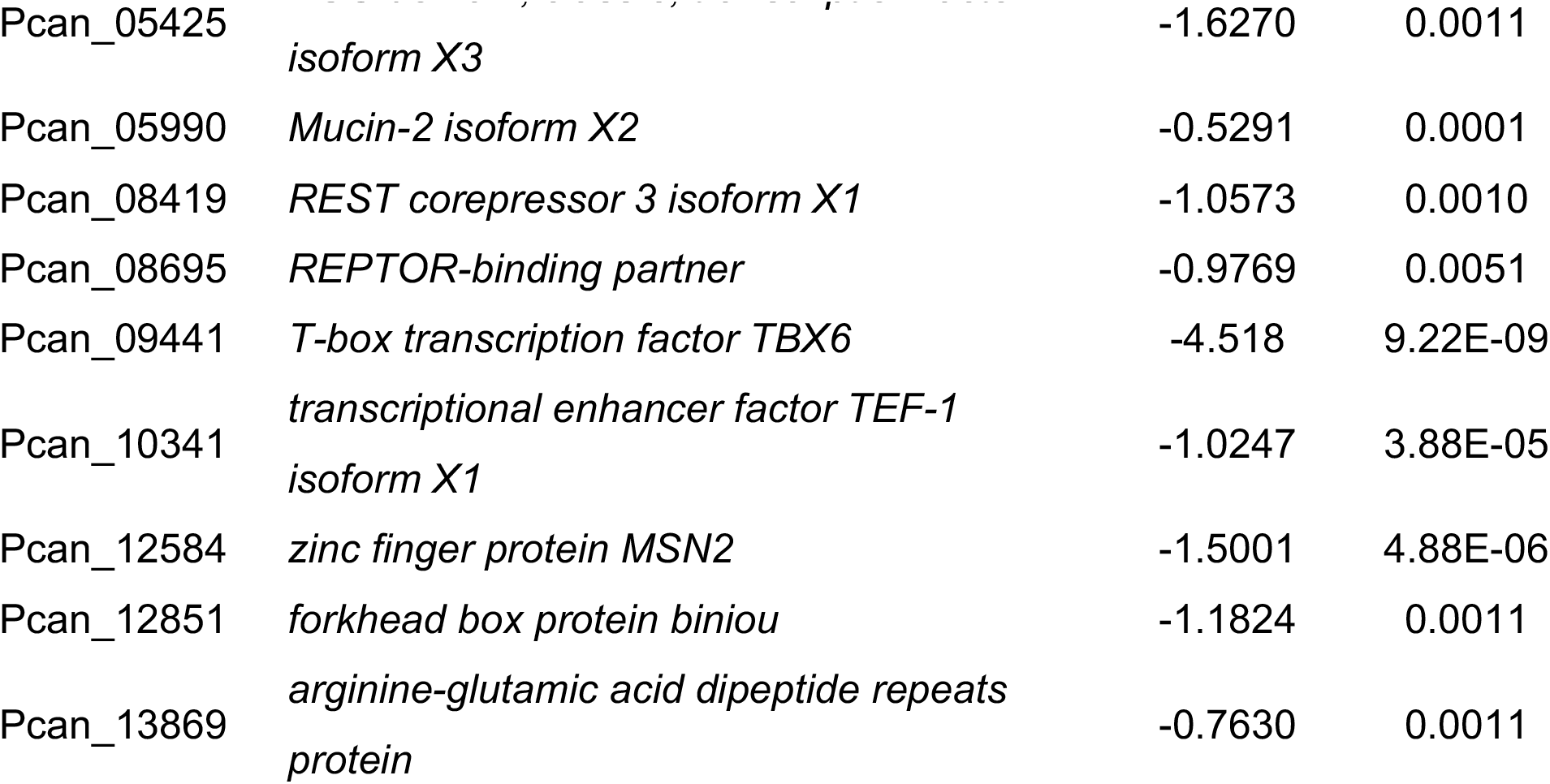
List of differentially expressed transcription factors. Their log2FC and padj values. The four genes (shaded in grey) at the top of the table are upregulated in spot (vs Cu1 samples), the remaining genes are downregulated in spots.

Recent studies have also pointed out the critical role that some signalling pathways have in organising butterfly wing patterns, providing positional information to surrounding cells to specify the development of colour patterns. Examples include but are not limited to Smad/TGF-β, Wnt, and the Hedgehog signalling pathway [29-31]. To infer the overall output of a signalling pathway, i.e, whether a pathway is active or repressed, expression profiles of all genes known to belong to signalling pathways in spot tissues were examined. In spots, the Toll, Notch, Transforming growth factor beta (TGF-β), Juvenile hormone and the Fibroblast growth factor (FGFR) pathways are likely to be associated with the development of spot patterns during early pupal stages (Table 4). Key positive regulators of these pathways were up-regulated in spots. In contrast, members of the Ecdysone and Hedgehog signalling pathway were down-regulated in spot tissues. The Wnt signalling pathway may also be repressed in spots (at 3-6 h), as key positive regulators of the Wnt pathway were down-regulated in spots (see discussion).

**Table 4.**
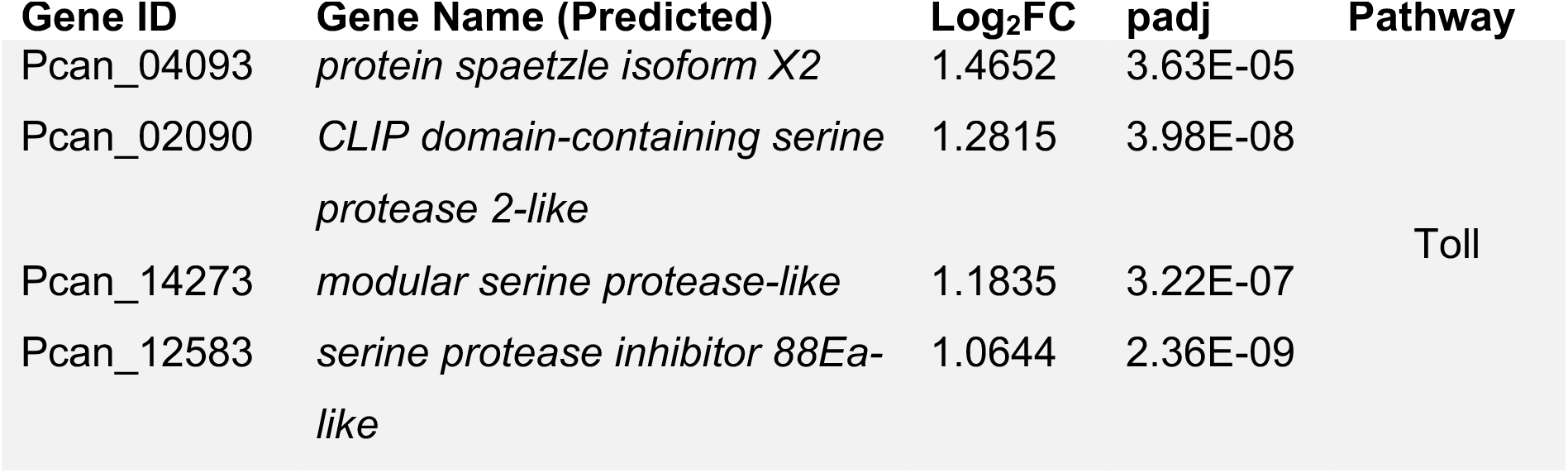
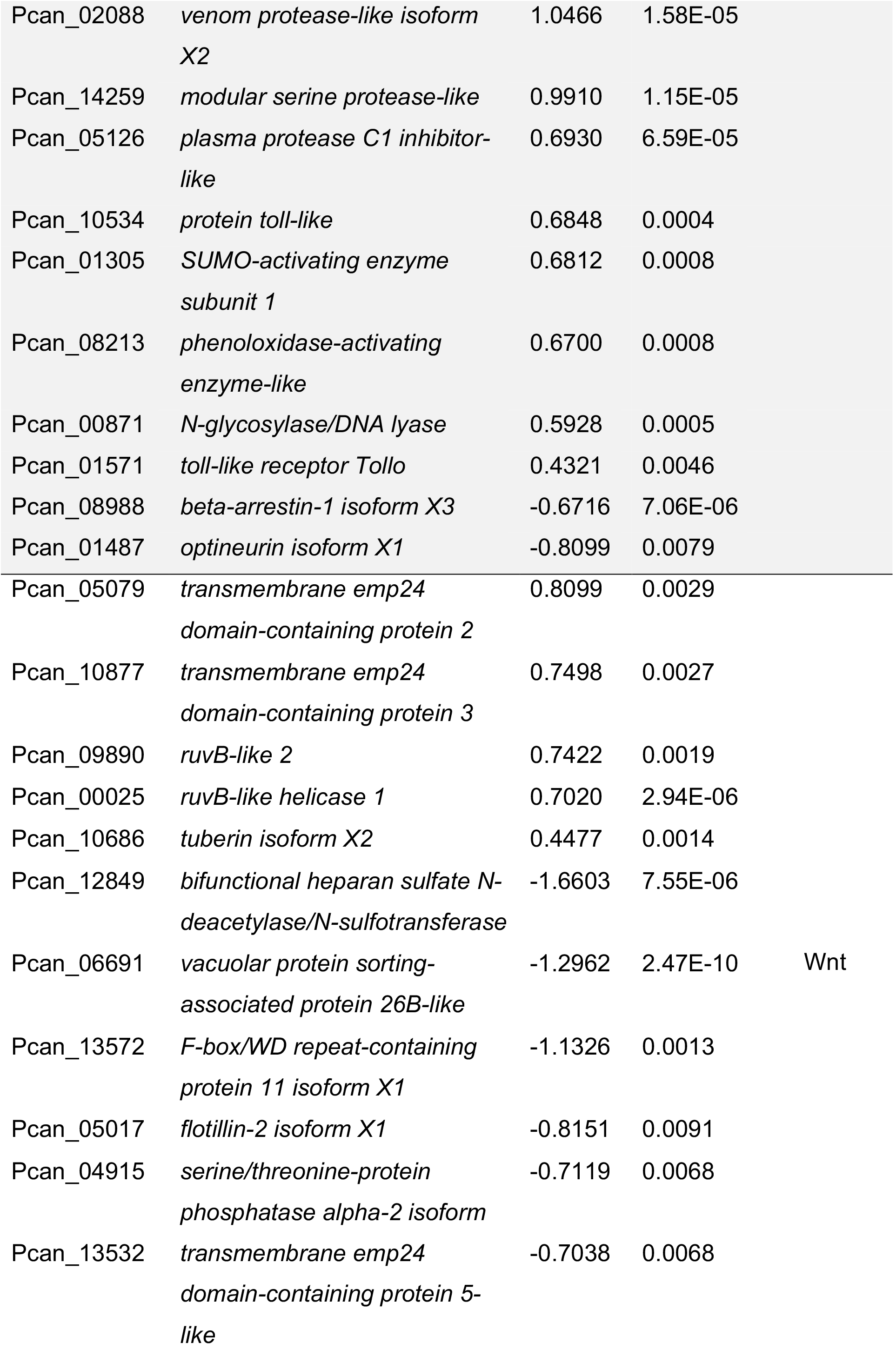
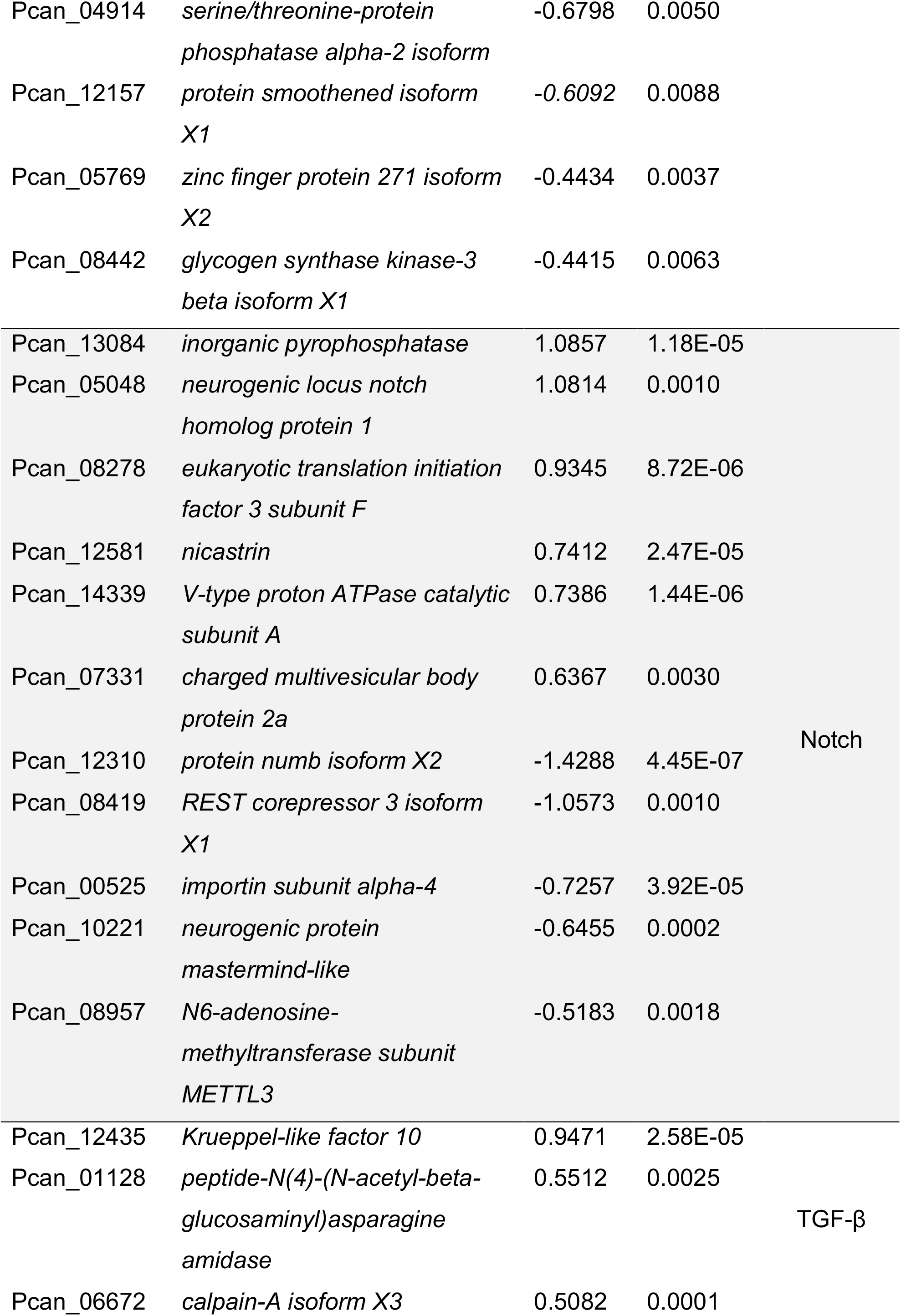
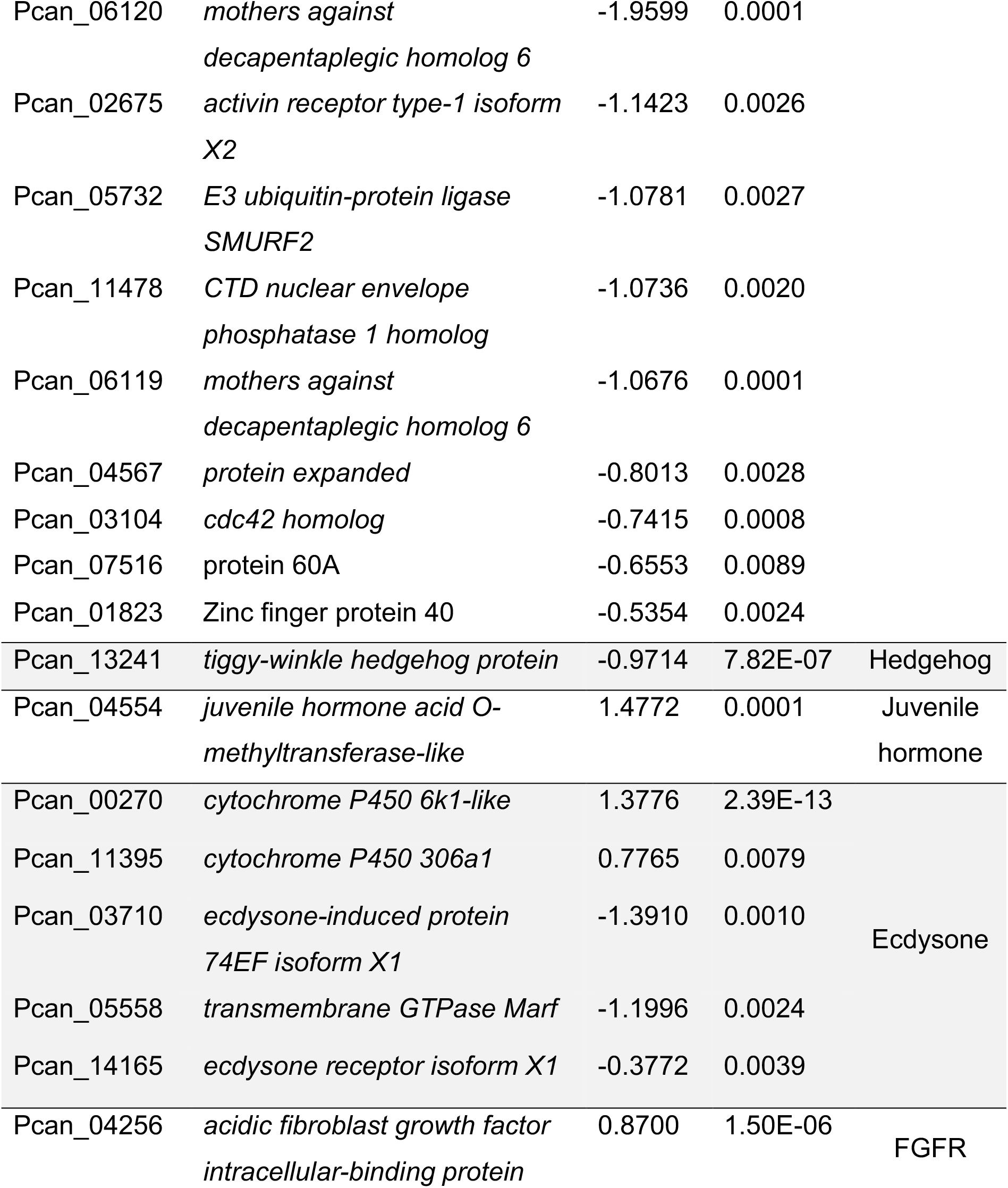
List of differentially expressed genes in Spot vs Cu1 tissues belonging to different signalling pathways. Their log2FC, padj values, and the signalling pathway they are associated with.

### Genes known to be functioning or expressed in eyespot development

To explore the possibility that eyespots and spots share common developmental modules, we manually cross-listed the DE genes obtained in this study with the list of genes known to be associated with eyespot development from published eyespot transcriptomes and the literature (Figure 3).

**Figure 3.**
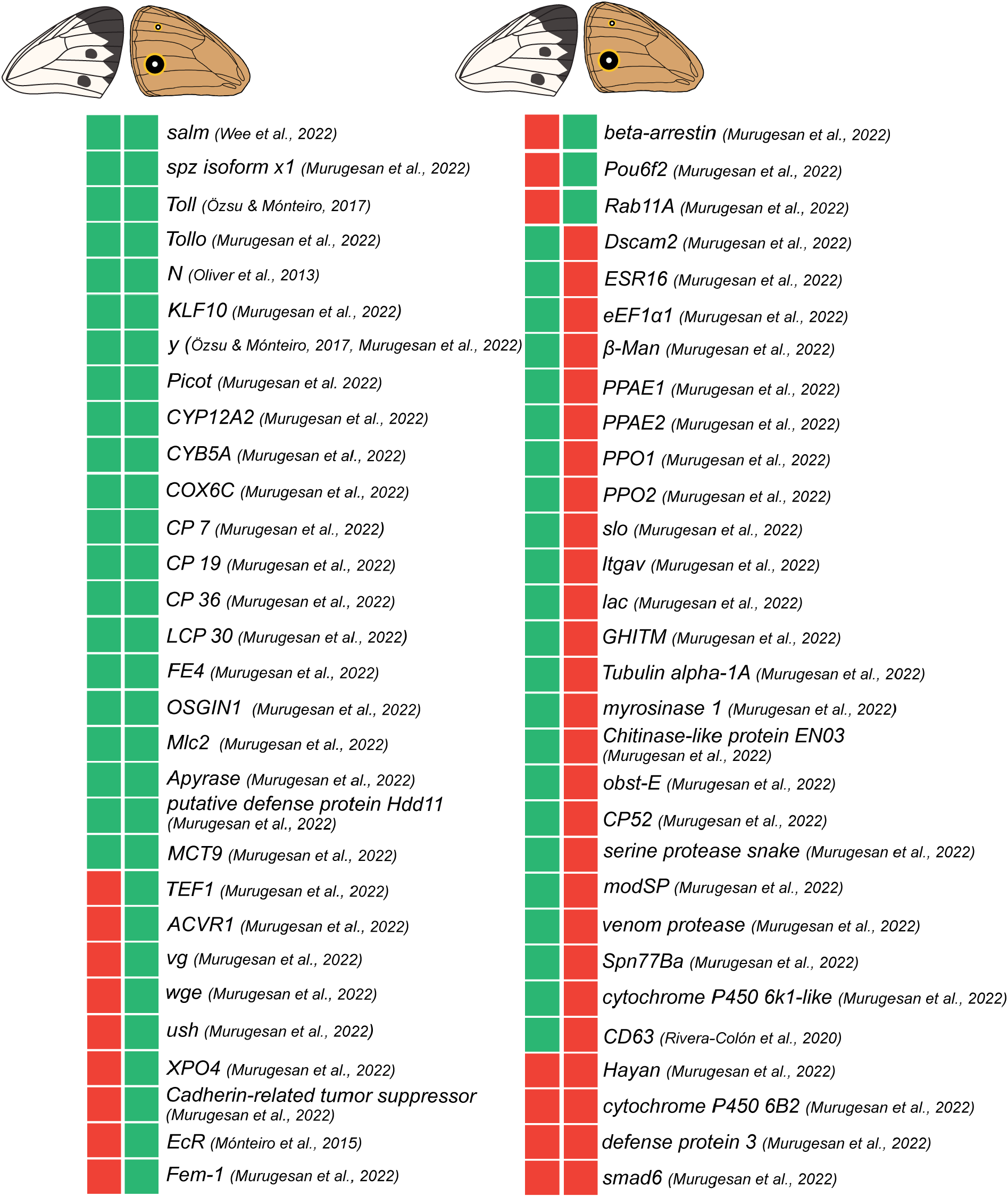
Core set of genes shared or DE between spot and eyespot patterns. DE genes in *P. canidia* spots and reference that describes the gene as also being DE in eyespots of *B. anynana*. Green blocks represent upregulated genes, while red blocks represent downregulated genes. Sum of expression patterns are: 21 genes are green-green, 12 genes are red-green, 23 genes are green-red and 4 genes are red-red.

## Discussion

Pioneers of pierid wing patterns research, such as Schwanwitsch [32] and Shapiro [33] proposed that spots of pierid butterflies are positional homologs of marginal nymphalid wing elements rather than of eyespots. This proposal was based on early wing pattern comparative work based on an extension of the nymphalid ground plan, a schematic of patterns homologies and symmetry systems deeply conserved amongst nymphalid species [34]. Subsequent studies on the molecular basis of these pierid wing spots showed that none of the known eyespot central (focal) genes were expressed in the centers of spots in both larval and pupal wings [10, 14, 35, 36]. The only exception was *spalt* which is expressed homogeneously in areas of the pupal wings that would eventually develop the complete black spot pattern. In addition to its role in differentiating eyespot centers, *spalt* is also an organiser of distal wing patterns in nymphalids. A robust expression of *Dpp*, an eyespot candidate morphogen, along the margins of *P. canidia* larval wings juxtaposed with the absence of *Dpp* signal in the center of spots [10], collectively supports the hypothesis that pierid spots may be variations of submarginal nymphalid pattern elements and not true homologs of nymphalid eyespot patterns. Here, however, we provide genetic evidence that the two pattern elements may have genes and developmental modules in common.

In this study we identified the first unbiased set of spot candidate genes in a pierid species at 3-6h post pupation. We discovered a large suite of pigmentation genes, transcription factors, and signalling pathways, differentially expressed in spots and flanking wing areas, that are likely working together to pre-pattern the development of a melanin spot. Interestingly, some of these genes are also expressed in eyespot centers, suggesting process homology at the level of GRNs between these two traits.

### Genes belonging to the melanin pathway are upregulated in spots

Most of the pigment-associated genes belong to the melanin biosynthesis pathway and were significantly upregulated in the presumptive spot area (Fig 4). These genes are described below.

**Figure 4.**
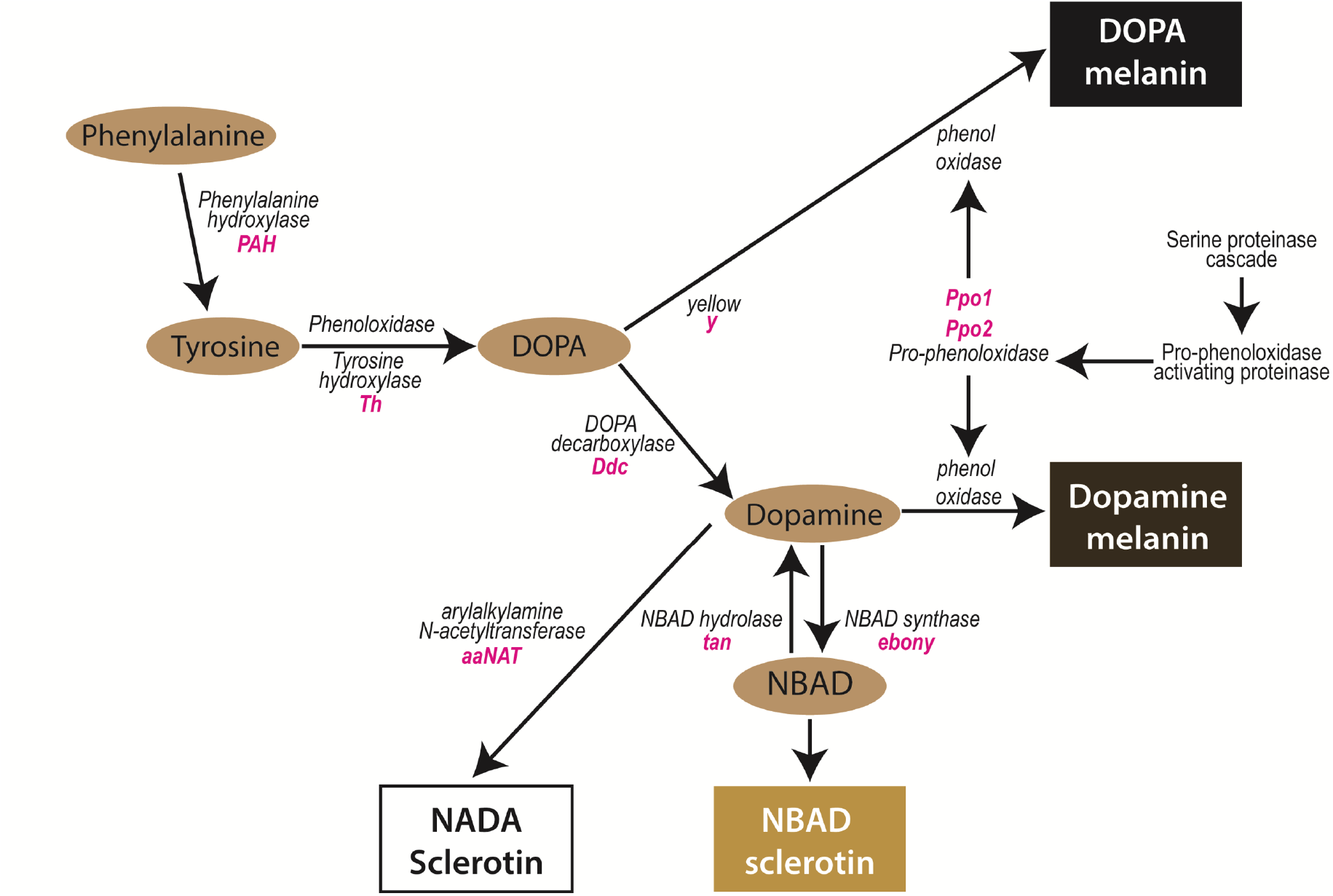
Part of the melanin biosynthesis pathway in insects. The enzymatic reactions that produce two types of melanins, DOPA melanin and dopamine melanin are shown here. Enzymes involved in the catalytic reactions of intermediate substrates are in black, and the genes that encode these enzymes are italicised in pink. Adapted from [37].

**Figure 5.**
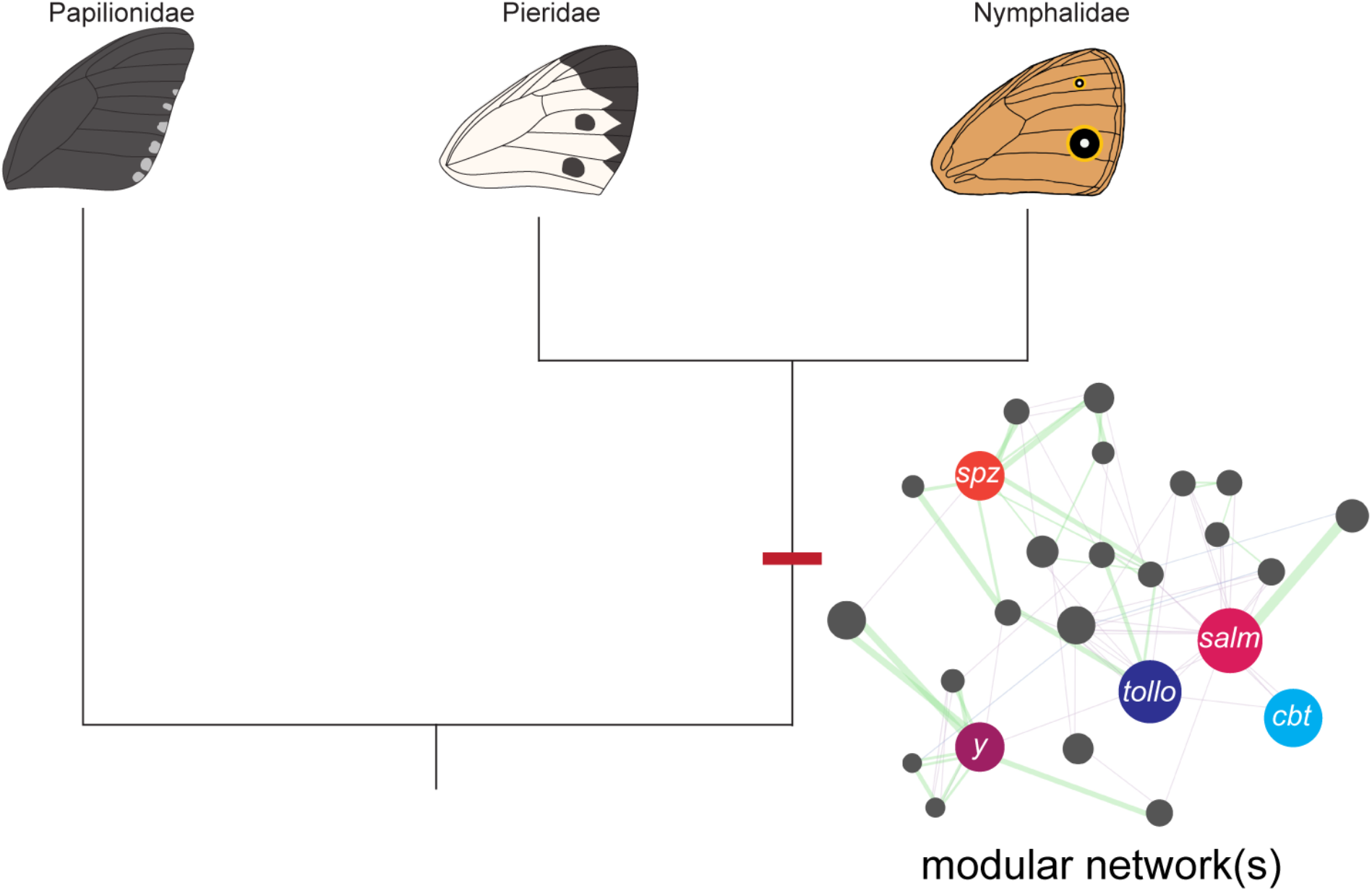
Eyespots and spots are non-homologous traits that might share homology at the level of GRNs. Both pierid spots and nymphalid eyespots may share common developmental modules. The core set of genes identified as common to both wing patterns may be a representation of an ancient ‘melanism’ module that was used in both patterns to produce melanin pigments or an ancient ‘wing patterning’ module that has been used repeatedly amongst butterfly species to organise the formation of patterns on the wing. However, more work needs to be done to validate the role(s) of this core set of genes in wing pattern development. Also, the function of gene enhancers common to these two non-homologous traits would also have to be studied to confirm that homology exists at the level of GRNs.

Several members of the *yellow* gene family, *yellow, L-dopachrome tautomerase yellow-f-like* and three copies of *yellow-like genes* were significantly up-regulated in wing spot tissues. Although the molecular functions of *yellow* genes are not yet fully determined, the expression of *yellow* prefigures melanic patterns and *yellow* genes are necessary for the formation of melanin pigments in *Drosophila, Bombyx mori*, numerous butterflies and many other insects [38]. *Yellow* was previously thought to have enzymatic properties that catalyses the conversion of melanin precursors into black melanin [39], but dopachrome conversion enzyme (DCE) activity was not detected in-vitro. Instead, a signal peptide that promotes the export of proteins is nestled within the N-terminus of the *yellow* protein coding sequence and Yellow protein has been visualised localising to extracellular regions. This suggests that *yellow* may regulate the formation of melanic patterns by acting as an anchoring protein that binds to melanin pigments in the cuticle [39-41]. However, in *Drosophila*, the two other related genes, *yellow-f* and *yellow f-2*, have DCE enzymatic activity so these genes would likely be directly involved in the synthesis of melanin pigments [42].

Like *yellow*, the two phenoloxidase (PO) enzymes, *phenoloxidase subunit 1*, and *phenoloxidase subunit 2*, that were upregulated in spot tissues, are also involved in the melanin pathway. POs are tyrosinases whose main role is to oxidise phenols to quinones, which polymerise to form melanin. In the melanin pathway, POs hydroxylate monophenols, such as tyrosine, to produce the melanin precursor DOPA.

Although part of the pteridine pathway, the catalytic product of *pyrimidodiazepine (PDA) synthase-like*, may contribute to the production of melanin pigments for spot development. *PDA synthase*, catalyses a reversible reaction of pyrimidodiazepine to 6-pyruvoyl-tetrahydropterin [43, 44]. 6-pyruvoyl-tetrahydropterin is a key intermediate required for the synthesis of tetrahydrobiopterin, also known as BH_4_ [45]. In insects, BH_4_ acts as an essential cofactor for other enzymes in several key reactions that involve the conversion of phenylalanine and tyrosine to melanin [46-48] (Figure 4). Thus, the upregulation of *PDA synthase* in *Pieris* spots suggests that part of the pteridine biosynthesis pathway is also used to produce melanin pigments.

Another pigment associated gene, *cysteine sulfinic acid decarboxylase*, was also upregulated in black spots. *Csad* was suggested to be the locus responsible for a melanic mutation in *B. anynana* larvae, known as *chocolate* [49]. This gene belongs to the pyridoxal phosphate (PLP) aspartate aminotransferase superfamily. PLPs serve as co-enzymes in many biochemical reactions, and one of its most well-studied family members is *dopa decarboxylase* (*ddc*), that catalyses the decarboxylation of L-dopa to produce dopamine [50]. Thus, it is highly likely that *csad* might also have direct enzymatic role on the melanin pathway.

Collectively, these genes are likely involved in the molecular processes required for the production melanin pigments for the wing spot pattern in *P. canidia*.

### Transcription factors regulating spot genes

Three transcription factors were significantly upregulated in spot tissues. They may be involved in the regulation of the pigmentation effector genes described above.

*Helix-loop-helix protein delilah-like* (*tx*), has been reported to be a marker gene that differentiates epidermal cells to facilitate muscle attachment in *Drosophila melanogaster* [51]. This gene is also important for specifying intervein regions, and adhesion of wing layers in *Drosophila. tx* is expressed in a homogenous fashion across the wing blade but is absent in the veins and is thought to be a mediator of several signalling pathways to specify the fate of intervein cells [52]. However, how this gene might be functioning in the context of the lepidopteran wing spot pattern is unclear.

*spalt* was also up-regulated in spot tissue in early pupal wings, which provided support for previous immuno-localization studies showing that *sal* expression correlates with melanic wing patterns in pierids [14, 36]. *sal*’s expression also supported functional studies, showing the essential role of this gene in the development of spots in *P. canidia* [10, 14]. However, it is still unclear how *spalt* is functionally connected to genes of the melanin pathway in this species.

Outside of pierids, however, *spalt* has also been associated with the development of both melanic and non-melanic patterns outside of pierids. In the pupal wings of the nymphalid *B. anynana, sal* expression maps to the black scales in eyespots and brown scales along the wing margin. This expression of *Sal* demarcating the boundaries of brown scales along the wing margin is also observed in another nymphalid species, *J. coenia* [53]. Additionally, in *J. coenia*, expression of *sal* is also associated with white and blue scales that are present within the eyespot pattern and in white spot patterns located on the forewing [53, 54] while in the lycaenid butterfly *L. melissa, sal* expression is associated with metallic-coloured scales located in chevrons at the margin [54]. These data suggest that *spalt* may play a primary pattern organisation role, rather than be a regulator of melanin pigmentation per se.

*Kruppel-like factor 10* (*Klf10*) belongs to the Specificity protein 1 (Sp1)/Krüppel-like zinc finger family of transcription factors and it was up-regulated in spots. Members of this family are characterised by three deeply conserved C2H2 zinc finger motifs that preferentially bind to GC-rich promoter regions to modulate transcription of target genes [55, 56]. Sp1/Krüppel-like transcription factors, also known as transforming growth factor-beta (TGF-β) inducible genes (TIEG), modulate numerous cellular processes such as epithelial cell proliferation, differentiation, apoptosis, circadian rhythm, and regeneration, among other processes [57-59]. In *Drosophila*, the fly homolog of *Klf10* is *cabut* (*cbt*). *cbt* is a positive regulator of TGF-β signalling and is known to be involved in wing disc morphogenesis, cell proliferation, and dorsal closure [60]. During *Drosophila* embryogenesis, *cbt* mutant embryos suffered from dorsal closure defects and expression of *dpp* was found to be downregulated in epidermal leading-edge cells during closure [61]. In *Drosophila* wing imaginal discs, *cbt* was found to be a positive regulator of *dpp* target genes such as *spalt* and *optomotor-blind* [60]. Although *dpp* was not detected in late larval wings, and 18h pupal wings in *P. canidia* [10], *dpp* might be specifying the spot pattern in early pupal wings (3-6h), either from the center of this pattern or from the wing margin [10]. *spalt* expression in spots may be responding to upstream signal input from both *dpp* and *cbt*.

### Pathways associated with spot development

The most highly represented signalling pathway in spot tissue was the Toll signalling pathway. Fourteen genes belonging or interacting with this pathway were upregulated in spots. The Toll pathway is involved in wound repair and facilitating the production of melanin at wound sites [62-64]. To activate the Toll pathway, the extracellular ligand, *spaeztle* (up-regulated in spots) must first be proteolytically cleaved by proteases such as *serine protease easter-like* (also up-regulated in spots) [65]. The cleaved ligand subsequently binds to the Toll receptor (up-regulated in spots) inducing a receptor dimerization process. The dimerised Toll receptor will then interact with a complex of several other downstream genes in a signalling cascade [66].

The Toll pathway has been previously implicated in the formation of wing patterns and melanic larval patterns in other lepidopteran species. Toll genes are overexpressed in eyespots of *B. anynana* [15], and in red marginal spots of *Papilio polytes* [67]. *Toll receptor 8* and *spätzle3* are required for the development of epidermal black stripes in the larvae of a silkworm mutant strain [68]. As mentioned previously, the melanin synthesis pathway is, in part, catalysed by the enzyme phenoloxidase (PO) which normally exists in its inactive form proPO. When required, proPO will be converted to active PO by a series of serine proteases, to eventually produce melanin pigments. In *Drosophila*, there is evidence for molecular cross-talks between the Toll pathway and melanin synthesis pathway as they are activated by a shared set of serine proteases working upstream of both pathways [69, 70] and it is likely that the melanin produced in *Pieris* black wing spots may have been synthesised through the PO-mediated melanin synthesis pathway.

Interestingly, the *Notch* receptor and other members of the Notch signalling pathway are overexpressed in spots. The Notch signalling pathway is known for its role in differentiating and patterning cells along the dorsal/ventral boundary that make up the wing margin in *Drosophila* [71-73]. One of the members of this pathway, *nicastrin*, shown here to be upregulated in spots, is an essential component for the activation of the pathway [74]. Other positive regulators expressed in spots include *eukaryotic translation initiation factor 3 subunit F* (*EIF3F*) and *V-type proton ATPase catalytic subunit A* (*Vha68-1*), which are required for the de-ubiquitination of the activated Notch receptor [75], and intracellular trafficking of Notch signals [76], respectively. *Notch* is expressed in midline patterns in *Pieris rapae* in late larval wing discs [35], but its function in butterfly wings is still unknown. Larval integument markings, however, have been previously disrupted via the knockdown of both *Notch* and *delta*, using siRNA, in three different species of lepidopteran larvae, *Papilio xuthus, Papilio machaon*, and the multi lunar (L) mutant strain of *Bombyx mori* [35, 77]. The Notch pathway may be patterning the wing spot in early pupal stages, but additional functional work is needed to functionally validate this hypothesis.

The Dpp/BMP (Bone morphogenetic proteins) branch of TGF-β signalling may also be upregulated in early pupal stages of spot development. Within the TGF-β pathway, there are two subfamilies of ligands, namely the BMPs and the activins [78]. These ligands, upon binding their receptors, use different sets of Smad genes, transcription factors that translocate to the nucleus, to transcriptionally modulate target genes [79]. Identification of the Smads being up-regulated in tissues allows the identification of which branch of the TGF-β signalling pathway is being used.

Within the list of DE genes, 2 copies of *mothers against decapentaplegic homolog 6 (Smad6)*, are downregulated in spot tissues. *Smad6*, whose fly homolog is *Daughters against dpp* (*Dad*), belongs to a class of inhibitory Smads whose function is to downregulate *Dpp* signalling [80, 81]. The downregulation of *Smad6* in spots, suggests that *dpp* signalling is active in spot tissues at 3-6h post pupation.

The up-regulation of *peptide-N(4)-(N-acetyl-beta-glucosaminyl) asparagine amidase*, known as *PngI*, also suggests that the *dpp* signalling is active in spot tissues at 3-6h post pupation, as this protein in known to upregulate *dpp* expression in the *Drosophila* gut [82].

*Wnt* genes have long been associated with the organisation and development of butterfly wing patterns [30, 83-85] but it is inconclusive as to whether the Wnt pathway is involved in spot formation in pierids. In *P. canidia*, two modulators of the Wnt pathway, *casein kinase II subunit beta* (*CK2*) and *bifunctional heparan sulfate N-deacetylase/N-sulfotransferase* (*sulfateless*) are downregulated in spots, but these modulators are not specific to the Wnt signalling pathway [86, 87]. Two other genes, *RuvB-like helicase 1* and *ruvB-like helicase 2*, are known antagonistic regulators of β-catenin activity, a signal transducer of Wnt signalling. *rvb1* is a positive regulator of Wnt signalling while the expression of *rvb2* represses Wnt/β-catenin signalling [88]. Since both genes are upregulated in spot tissues, they provide limited evidence as to whether Wnt signalling is active in spots. Interestingly, *frizzled-2*, the receptor for *wingless* [89, 90], was down-regulated in spot-containing tissue, albeit at a higher padj value of 0.013 and with a Log_2_FC of -0.64. Since Wnt signalling is known to repress the expression of *frizzled-2* on the *Drosophila* wing to shape and sustain the Wg morphogen gradient [91], future work should study the role of Wg signalling in organising and positioning pierid spot patterns along the margin.

Our data also points to the insect hormone JH in the regulation of spot size. JH is one of the two major insect hormones that regulate the development of seasonal polyphenisms [92, 93], and *P. canidia* spot patterns are seasonally plastic. In colder seasons, these wing spots are bigger and darker [94]. Our data shows that the gene *juvenile hormone acid O-methyltransferase-like*, is upregulated in spot tissues. This gene encodes an enzyme that mediates the conversion of inactive precursors to JH [95]. Future work attempting to dissect the seasonal plasticity of *P. canidia* wings should look to studying the possible functions of JH in regulating the size and intensity of the melanised spots between the two seasonal forms.

Spots and eyespots are non-homologous wing pattern elements that may share developmental modules in common. We compared the spot transcriptome of *P. canidia* to two different eyespot transcriptomes from *B. anynana*, all sampled at 3-6h after pupation [7, 15]. This current spot transcriptome was also cross checked with previous studies of genes associated with eyespot development. Collectively, these comparisons revealed a set of shared genes that are expressed in both traits (Figure 4).

Genes and signalling pathways shared across the two non-homologous traits include *spalt, yellow, notch*, Toll, and BMP signalling. Some of these genes (*yellow, spalt*, and Toll genes) may simply reflect the independent deployment of an ancient GRN, that was deeply conserved across numerous taxa (i.e., melanin gene regulatory network), onto the wing to produce black pigments needed to form the black circles in both spots and eyespots. Other genes and pathways such as *(Klf-10, Notch*, FGFR and TGF-β) may likely also have conserved roles in organising wing patterns along the tip of the midline of each wing sector in insects. It remains to be seen, however, whether genes shared between eyespots and spots, and their regulatory interactions, were recruited once, in the ancestors of both lineages, or evolved convergently to function in wing patterning, independently in each lineage. To help solve this question, a more comprehensive analysis of spot evolution is necessary, using a denser sampling of species across all butterflies. Alternatively, discovering that genes common to spots and eyespots sharing the same enhancers might also confirm process homology across both wing patterns.

### Conclusion

Here, we report a complete genome assembly of *Pieris canidia* and identify transcripts associated with early wing spot formation in this species. The list of DEGs identified here, represents the first unbiased set of spot candidate genes in a pierid species and provides a comparative framework to study the origin and diversification of butterfly wing patterns. These analyses, however, represents a single snapshot of spot development during a specific developmental stage. Future studies should functionally validate the list of identified spot genes using CRISPR/Cas9 system for further insights on the developmental basis of spot patterns.

## Materials and Methods

### Animal Husbandry

*Pieris canidia* used in this study were filial descendants from wild-caught individuals found along West Seletar Road, Singapore. Caterpillars were reared on potted *Brassica sp*. plants, at 27°C, 80% humidity following a 12h light:12h dark cycle. *P. canidia* pre-pupae are readily identified by several changes to their morphology and behaviour. During the last stage of larval development, *P. canidia* larvae will shrink slightly in length and move away from the potted plants. Once a suitable substrate for pupation is found, the larvae will produce a silk girdle around their thoracic segments and attach to the substrate. These pre-pupae were carefully removed from their pupation site and transferred to petri dishes where their time of pupation was monitored using a camera (Olympus Stylus Tough TG-5 Digital Camera) with the in-built time-lapse function. Pupae that were 3-6h old were sexed, with only female pupae used for subsequent dissections to exclude any gender bias in gene expression profiles as *P. canidia* wing spots are sexually dimorphic. Female spots are larger than male spots.

### Genome Assembly

High Molecular weight (HMW) DNA was extracted from two female pupae using a phenol-chloroform method with little modification [96]. Extracted DNA was stored in - 80°C before being pooled and sent for sequencing. Long reads library preparation and sequencing was performed at AIT Novogene, Singapore. 20GB of data, roughly of about (70x) coverage was sequenced using the PacBio sequel platform.

Initial genome assembly was carried out using canu [19] and wtdbg2 [18] assemblers separately. The canu assembler was run with the default option with minimum read length set to 3000 for the assembly, and wtdbg2 was run with default settings. For both assemblies, the relative genome size of 320 MB was set based on the sister species genome sizes (*P. napi* and *P. rapae*) [97]. The two assemblies were merged using quickmerge [20], with the canu assembly as the reference, and the wtdbg2 assembly as the query. Heterozygosity in the contigs of the resulting assembly was purged using purge_haplotigs [21], and only the haploid contigs were kept. The obtained contigs were polished with three rounds of racon [22] and Pilon [23] using corrected PacBio long reads and Illumina short reads, respectively, to get the final genome assembly, Pcan.v1. Illumina short reads were generated using standard protocol. DNA was extracted from a female adult individual. Library preparation was done using the Illumina TruSeq Nano DNA kit, with cluster amplification performed using the Illumina cBot. Sequencing was done through the Illumina HiSeqX platform (HiSeq Control Software 3.3.76/RTA 2.7.6) using sequencing-by-synthesis (SBS) kits to generate 151 bp paired-end reads. A BUSCO score [26] was used to check the completeness of the gene sets in the assembly resulting in 14518 genes with 26376 transcripts.

### Repeat Masking of Genome

The genome was repeat masked for transposable elements, small and tandem repeats. Repeat Masking was performed using a similar approach as that used for *B. anynana* genome masking [7]. De novo transposable elements and repeats were modelled and predicted using RepeatModeler v1.0.5 [98]. The repeat library created using the *de novo* approach sometimes had non-repeats and transposable elements in them, and to filter the non-repeats and coding regions we used the protein sequences from the genome annotation performed earlier. The resulting library was merged with the lepidoptera repeat library obtained from Rephase v20.05 [98]. The genome was masked using the final merged library using Repeatmasker v4.0.7 [99] resulting in 30.05 % of the genome being masked.

### Genome Annotation

The soft-repeatmasked genome was annotated, using four rounds of Maker v3 [25]. The transcriptome assembled from the RNA-Seq data was used as transcripts for the species, with transcriptome and protein sequences from *Pieris napi, Pieris rapae, Junonia coenia, Bombyx mori*, and *Bicyclus anynana* as relative species transcripts and protein homology evidence for the first round of gene predictions. Output gene predictions from each round were used as input for the next round. Snap and Augustus [100] were used for the second round of gene predictions, followed by Genemark [101] for the third round of gene modelling, and one final round of Snap and Augustus predictions. The minimum length of 35 amino acids was set for gene predictions. The predicted gene models were kept for genes that had an Annotation Edit Distance (AED) score of < 1 and/or had a gene ontology obtained from Interproscan [102]. This resulted in 14594 genes with 26,463 transcripts. In order to correct the annotations and produce a standardized gff3 file, the gff file obtained from Maker was run through agat_convert_sp_gxf2gxf.pl script, which is a part of AGAT tools [103]. This step resulted in the removal of 77 identical isoforms and added the missing gene features, leading to a total of 14518 genes with 26,376 transcripts. Functional annotation was performed by locally blasting the transcripts against a non-redundant (nr) protein database, using diamond blast [104], and a gene ontology analysis was performed using Interproscan in Blast2Go [105]. Finally, the blast results were merged with the Interproscan results in Blast2Go to produce a final functional annotation for the genome.

### Tissue collection and RNA extraction

Small squares of wing tissues of approximately 0.5 × 0.5 mm containing the spot centers from the M3 and Cu1 sector of the pupa were manually dissected from the dorsal forewing using a homemade dissecting tool which consists of thin strips of cut razor blades glued to a wooden handle (Fig S4). All dissections were made in cold diethyl pyrocarbonate (DEPC) treated 1xPBS and dissecting tools were wiped with RNaseZap® (Invitrogen™) prior to dissection. Dissections of the wing tissues of different wing sectors were done in a random fashion to avoid any bias in gene expression levels. Harvested wing tissues from the pupa were immediately placed into individual 1.5ml eppendorf tubes containing RNAlater® (Ambion) to preserve the integrity of the RNA. These tubes were then stored at 4°C for at least 24h to allow for complete RNA stabilisation and then transferred to -80°C storage until all samples required for four biological replicates were obtained. Around 30 pieces of tissue belonging to the same wing sector were pooled together, snap-frozen in liquid nitrogen, and homogenised using disposable polypropylene pellet pestles. RNA was subsequently extracted using the RNeasy Mini Kit (Qiagen), according to manufacturer’s instructions. Preliminary concentration of the extracted RNA was measured using a NanoDrop 100 Spectrophotometer (ND-1000) and RNA integrity was assessed through agarose gel electrophoresis and Agilent 2100 Bioanalyser by an external vendor (AITbiotech).

### Next generation sequencing

Library preparation, and sequencing was performed by Novogene. A total of 12 libraries was sequenced across all three different groups of wing tissues at 150-bp paired end using the Illumina NovaSeq6000 platform (Table S3.1 & S3.2).

### Pre-processing of raw sequencing data

Quality control, removal of adaptors, and read filtering was performed using *fastp* [106]. As the Novaseq sequencing platform was used to sequence the libraries, we chose *fastp* to remove overrepresented polyG tails that might appear from a lack of signal in a two-colour system used in Novaseq sequencing technologies.

### Genome alignment

A genome-guided transcriptome assembly was constructed using the HISAT2/StringTie pipeline [107] against the assembled *Pieris* genome (Table S3). Filtered reads were mapped to the *P. canidia* genome using HISAT2. The resulting alignments were later processed and quantified using StringTie which produces estimates of transcript abundances and generated read count tables. A python script prepDe.py was used to convert the output files from Stringtie to produce genes and transcripts count matrices in the form of CSV files.

### Differential gene expression analysis

Differential expression of genes was calculated using the Bioconductor R package, DESeq2 [108]. Pre-filtering of the gene count matrix was conducted prior to downstream analyses. Genes that had read counts present in less than three samples, and with normalised counts lesser or equal to 10 were removed from the dataset. A pairwise DE analysis was then made between the M3 spot tissue and the Cu1 non-spot control tissue. Genes with an adjusted P value of < 0.01 were considered as significantly differentially expressed in this dataset. In all, a total of three biological replicates for two different tissue groups (Spot & Cu1) were used in the final analysis.

### Annotation and GO annotation of DE genes

Differentially expressed genes were blasted against the nr database in NCBI using blastx in Blast2GO with an *E*-value of < 1e-3 with the application of a taxonomy filter Blast2GO and against both the InterPro and Eggnog databases. All GO annotations obtained through the three different databases for each transcript were then merged in the Blast2GO pipeline [105].

## Acknowledgements

We would like to thank Heidi Connahs for providing initial comments for the manuscript.

## Funding

This work was funded by the National Research Foundation (NRF) Singapore, under its Investigatorship Programme (NRF-NRFI05-2019-0006 Award). JW and SNM were also supported by graduate scholarships awarded by the Department of Biological Sciences, National University of Singapore, and by Yale-NUS, Singapore respectively.

## Availability of data and materials

All data used in the analyses are included within the article and its accompanying supplementary files. Raw Illumina and PacBio sequence reads used for the assembly of the genome and for the transcriptomic analyses are available under NCBI Bioproject PRJNA895987.

## Supplementary Materials

**Table S1.**
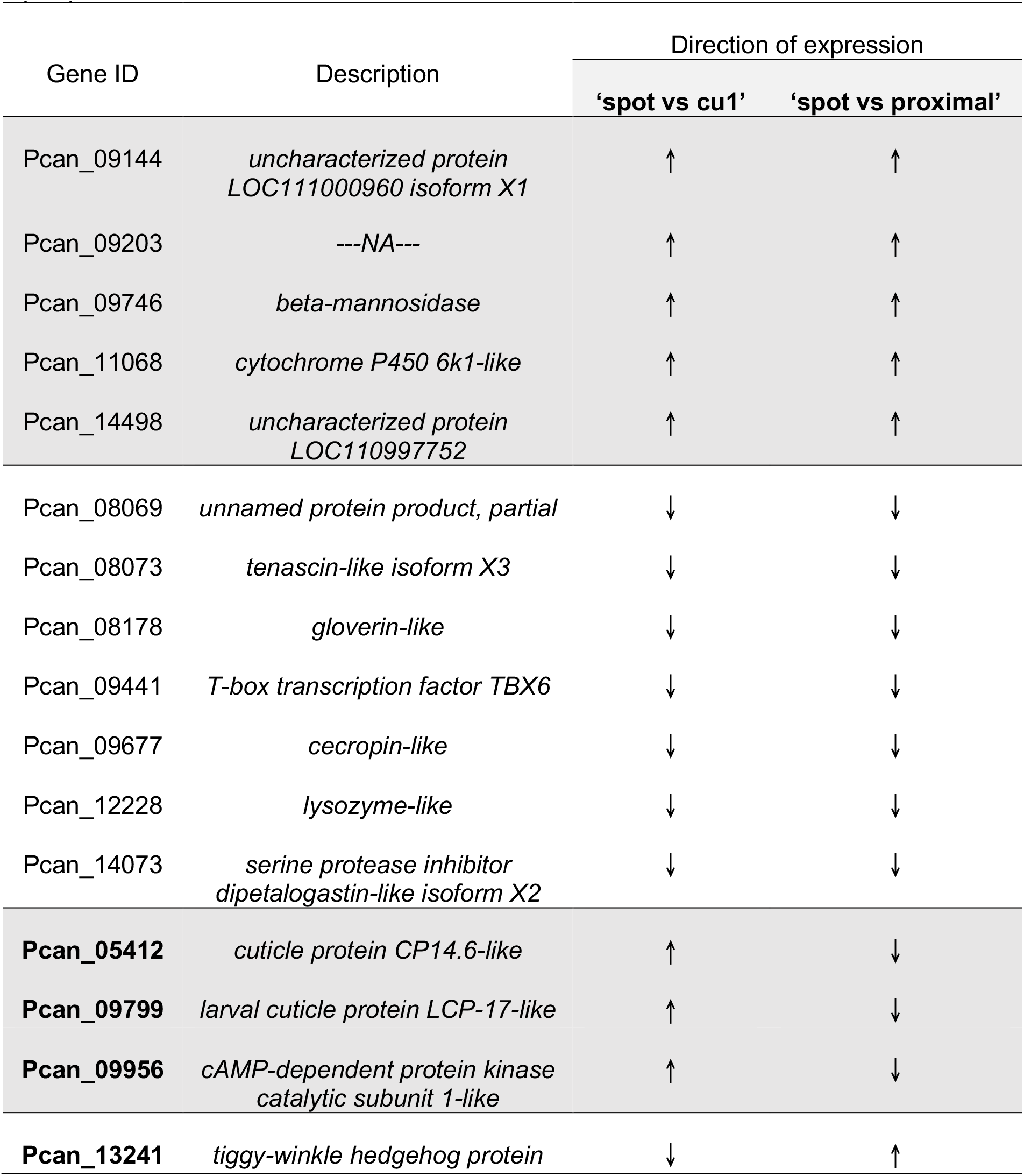
List of differentially regulated genes common to both set of comparisons: i) spot vs cu1 and ii) spot vs proximal. Up arrow symbol represents the upregulation of genes in wing tissues containing the ‘spot pattern’ while the down arrow symbol represents the downregulation of genes in wing tissues containing the ‘spot pattern’.

| Gene ID | Description | Direction of expression |  |
| --- | --- | --- | --- |
|  |  | 'spot vs cu1' | 'spot vs proximal' |
| Pcan_09144 | *uncharacterized protein LOC111000960 isoform X1* | ↑ | ↑ |
| Pcan_09203 | ---NA--- | ↑ | ↑ |
| Pcan_09746 | *beta-mannosidase* | ↑ | ↑ |
| Pcan_11068 | *cytochrome P450 6k1-like* | ↑ | ↑ |
| Pcan_14498 | *uncharacterized protein LOC110997752* | ↑ | ↑ |
| Pcan_08069 | *unnamed protein product, partial* | ↓ | ↓ |
| Pcan_08073 | *tenascin-like isoform X3* | ↓ | ↓ |
| Pcan_08178 | *gloverin-like* | ↓ | ↓ |
| Pcan_09441 | *T-box transcription factor TBX6* | ↓ | ↓ |
| Pcan_09677 | *cecropin-like* | ↓ | ↓ |
| Pcan_12228 | *lysozyme-like* | ↓ | ↓ |
| Pcan_14073 | *serine protease inhibitor dipetalogastin-like isoform X2* | ↓ | ↓ |
| **Pcan_05412** | *cuticle protein CP14.6-like* | ↑ | ↓ |
| **Pcan_09799** | *larval cuticle protein LCP-17-like* | ↑ | ↓ |
| **Pcan_09956** | *cAMP-dependent protein kinase catalytic subunit 1-like* | ↑ | ↓ |
| **Pcan_13241** | *tiggy-winkle hedgehog protein* | ↓ | ↑ |

**Table S2.** RNA concentrations obtained from dissected tissues for the 12 different libraries.

| Sample Name | Treatment Group | Concentration (ng/μl) | Total Volume |
| --- | --- | --- | --- |
| S1 | Spot | 17.6 | 39 |
| S2 | Spot | 32.7 | 42 |
| S3 | Spot | 12.8 | 42 |
| S4 | Spot | 25.2 | 60 |
| P1 | Proximal | 13.1 | 42 |
| P2 | Proximal | 16.7 | 42 |
| P3 | Proximal | 21.6 | 42 |
| P4 | Proximal | 21.7 | 60 |
| C1 | Cu1 | 33.8 | 42 |
| C2 | Cu1 | 27.1 | 60 |
| C3 | Cu1 | 32.3 | 60 |
| C4 | Cu1 | 21.4 | 60 |

**Table S3.** Summary of sequencing statistics and percentage of reads mapped to *P. canidia* genome using HISAT2.

| Sample | Raw Reads | Effective Rate | Q20 (%) | Q30 (%) | GC Content (%) | Percentage of filtered reads mapping to genome (%) |
| --- | --- | --- | --- | --- | --- | --- |
| **S1** | 54707539 | 97.15 | 98.06 | 94.19 | 50.04 | 96.40 |
| **S2** | 62825665 | 97.21 | 98.21 | 94.53 | 51.38 | 92.71 |
| **S3** | 48529341 | 97.25 | 98.16 | 94.39 | 51.48 | 96.98 |
| **S4** | 95642456 | 98.27 | 98.32 | 94.91 | 47.30 | 96.91 |
| **P1** | 51952490 | 97.88 | 98.25 | 94.63 | 50.39 | 96.70 |
| **P2** | 50864060 | 98.80 | 97.97 | 93.94 | 49.97 | 96.20 |
| **P3** | 48023578 | 97.89 | 98.20 | 94.47 | 50.59 | 96.88 |
| **P4** | 94616878 | 98.48 | 98.03 | 94.29 | 47.46 | 96.64 |
| **C1** | 61010193 | 97.83 | 98.09 | 94.28 | 51.40 | 96.85 |
| **C2** | 86241796 | 98.76 | 97.92 | 94.08 | 48.01 | 96.72 |
| **C3** | 98461684 | 98.52 | 98.07 | 94.42 | 49.04 | 96.74 |
| **C4** | 98364038 | 98.21 | 98.47 | 95.29 | 48.12 | 96.82 |

**Figure S1.**
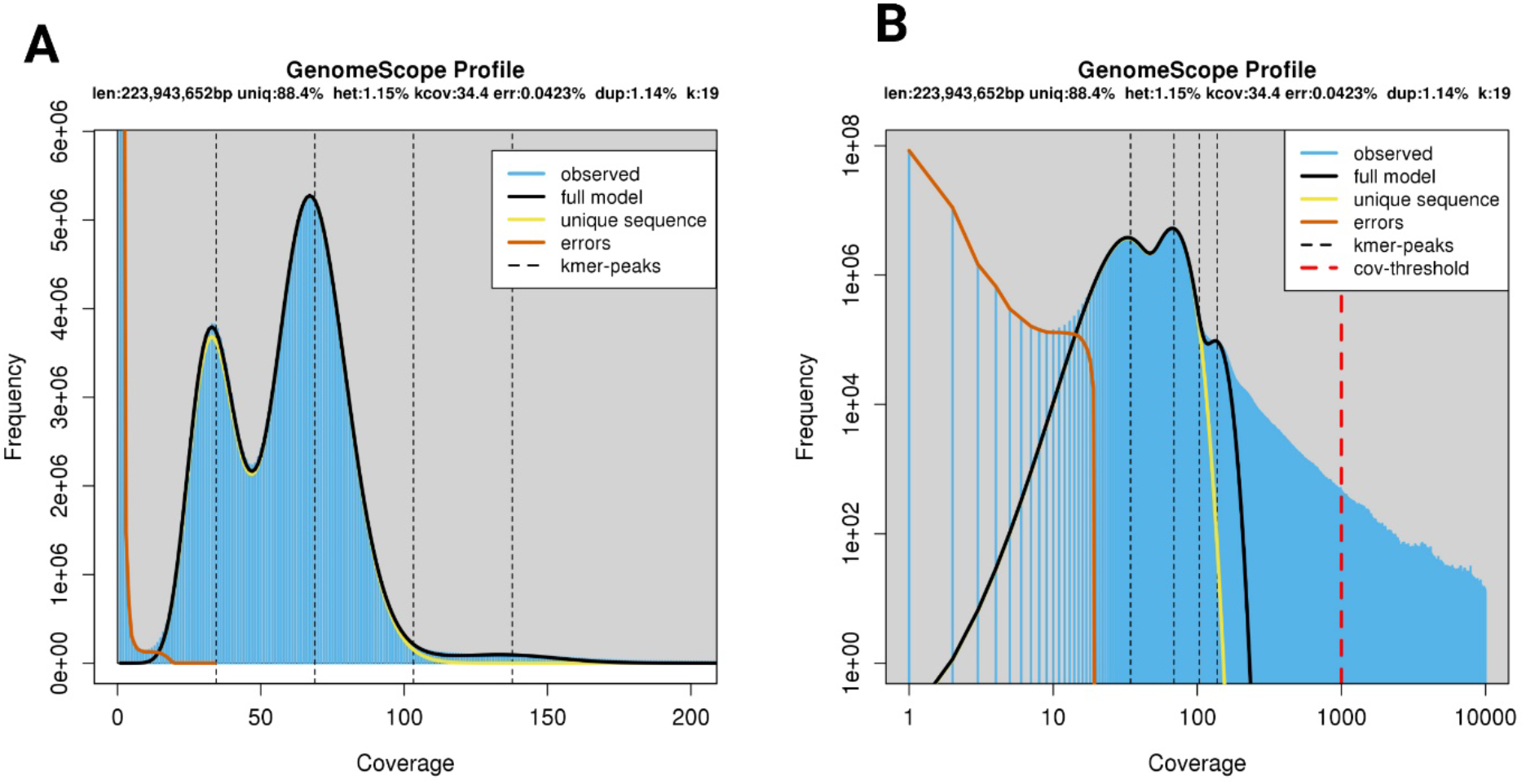
*P. canidia* genome size estimation using GenomeScope. We used Illumina short reads to estimate the genome size. We input the k-mer (k=19) analysis output from Jellyfish (Marçais & Kingsford, 2011) to estimate the haploid genome size. (A) Linear and (B) log plot of a k-mer spectral genome composition from *P. canidia* Illumina short read library. K-mer=19, Max K-mer coverage=1000. The estimated genome size was around 224 MB.

**Figure S2.**
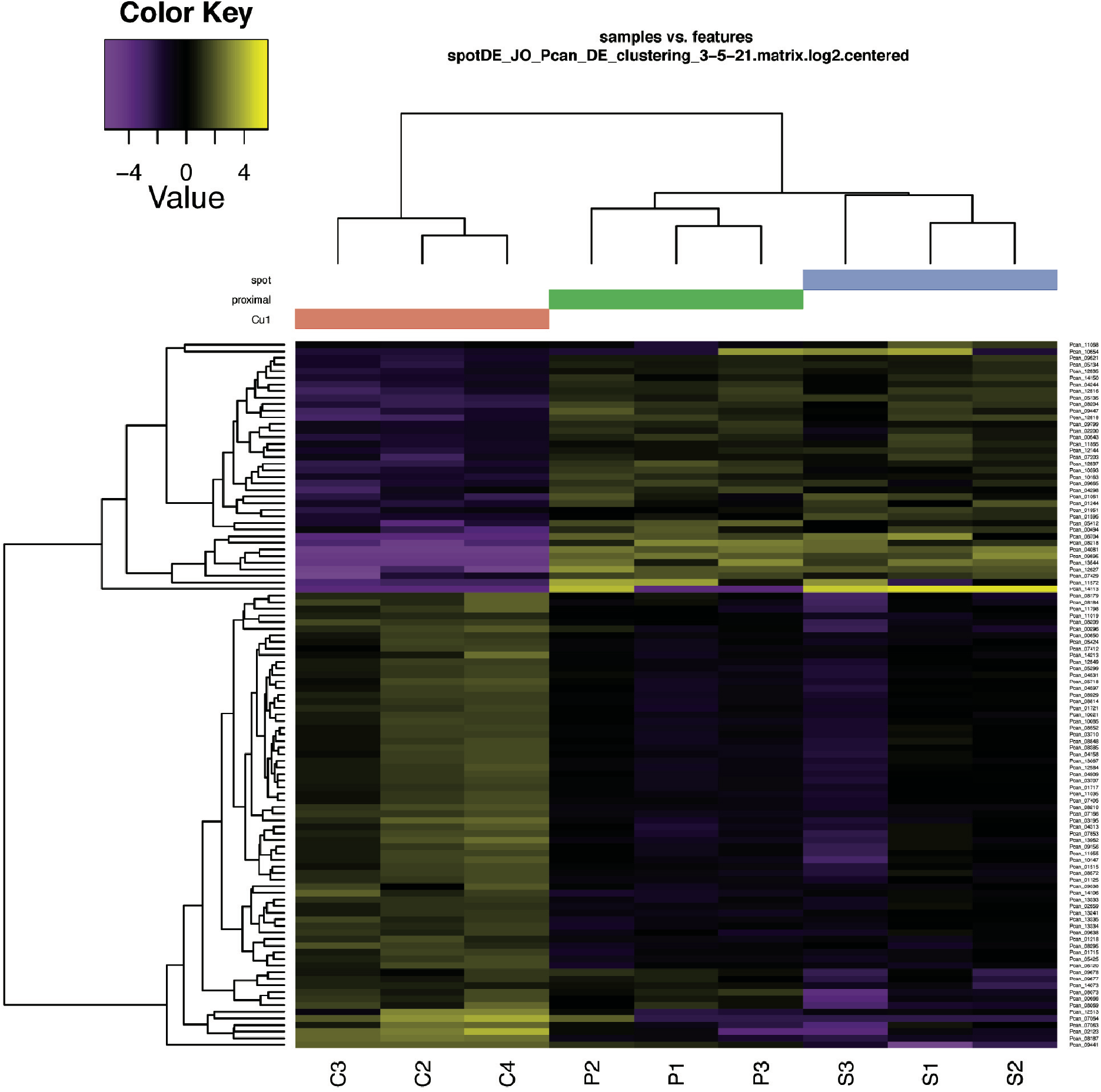
Hierarchical clustering and heat map. This clustering is based on the expression profiles of DE genes between “spot”, “proximal” and “cu1” groups for a set of significant genes (Padj <0.01).

**Figure S3.**
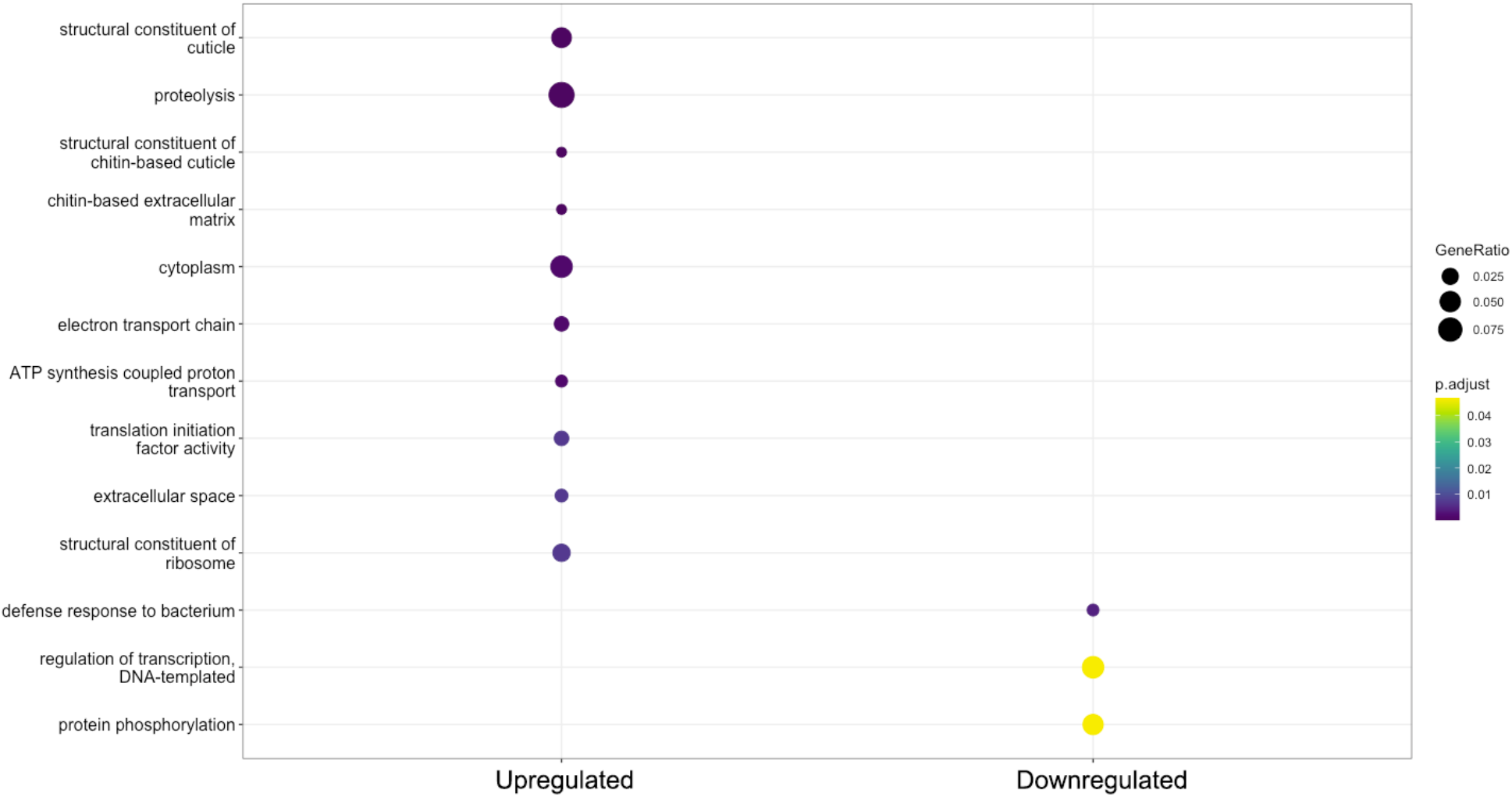
Gene Ontology (GO) enrichment analyses for spot DEGs. The function ‘*compareCluster’* within the Bioconductor package clusterProfiler 4.0 was used to determine over-represented GO terms in upregulated and downregulated DEGs, with the size of the dot denoting gene ratio and the colour of the dots representing the significance. P-values are adjusted by the Benjamini-Hochberg (BH) method.

**Figure S4.**
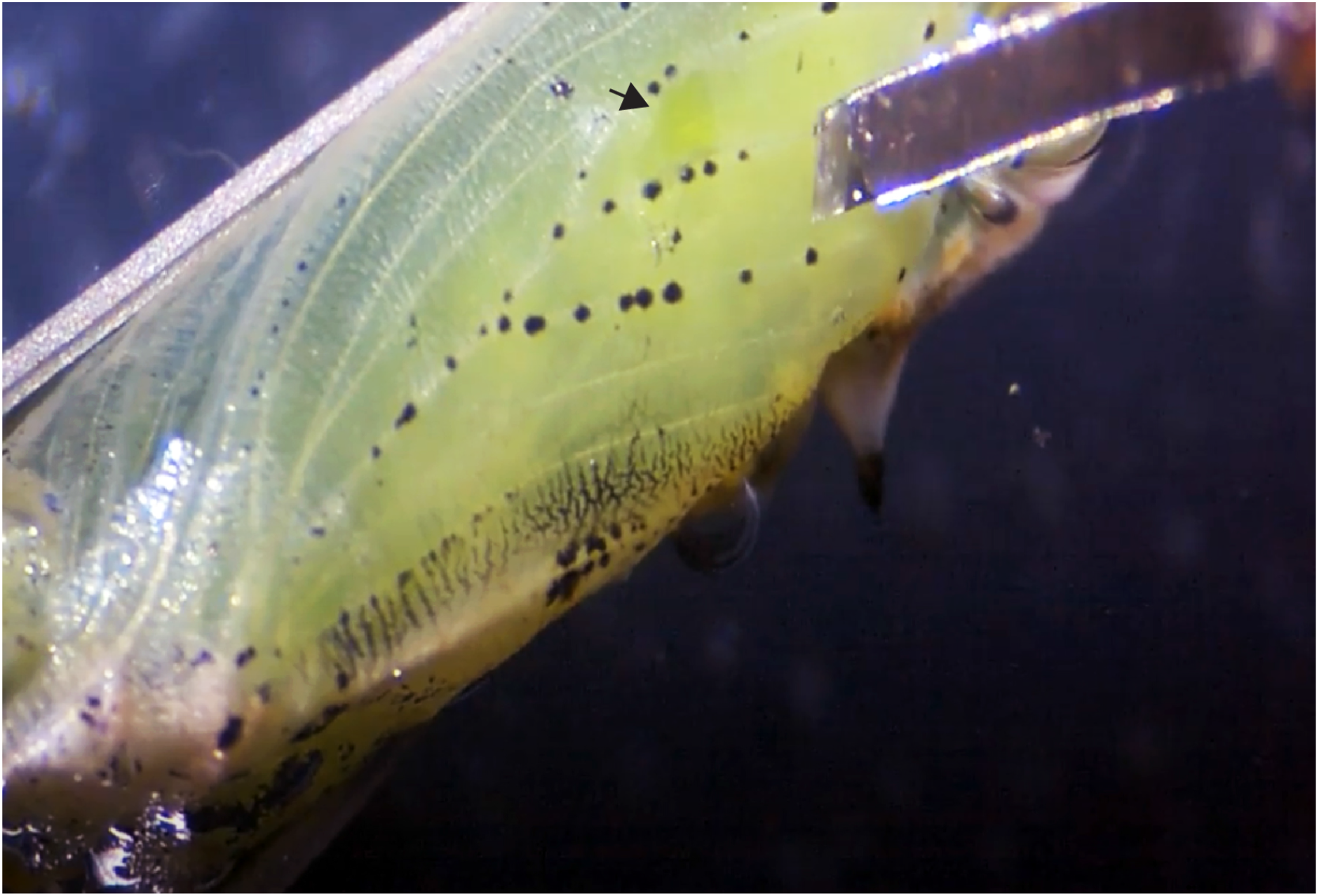
Microdissections of early pupal wing tissues using dissecting tools fashioned from cut razor blades. The black arrow denotes a piece of tissue that was already removed from the M3 wing sector.

## Notes

### Competing Interest Statement

The authors have declared no competing interest.

